# First description of *Lotmaria passim* and *Crithidia mellificae* haptomonad stage in the honeybee hindgut

**DOI:** 10.1101/2021.04.12.439428

**Authors:** María Buendía-Abad, Pilar García-Palencia, Luis Miguel de Pablos-Torró, José María Alunda, Antonio Osuna, Raquel Martín-Hernández, Mariano Higes

## Abstract

The remodelling of flagella into attachment structures is a common and important event in the insect stages of the trypanosomatid life cycle. Among their hymenopteran hosts, *Lotmaria passim* and *Crithidia mellificae* can parasitize *Apis mellifera*, and as a result they might have a significant impact on honeybee health. However, there are details of their life cycle and the mechanisms underlying their pathogenicity in this host that remain unclear. Here we show that both *L. passim* promastigotes and *C. mellificae* choanomastigotes differentiate into haptomonad stage covering the ileum and rectum of honeybees. These haptomonad cells remain attached to the host surface via zonular hemidesmosome-like structures, as revealed by Transmission Electron Microscopy. Hence, for the first time this work describes the haptomonad morphotype of these species and their hemidesmosome-like attachment in *Apis mellifera*, a key trait exploited by other trypanosomatid species to proliferate in the insect host hindgut.

**Author summary:** In recent years, the mortality of European Honeybees (*Apis mellifera*) has risen worldwide due to a variety of factors, including their infection by parasites. Former studies have linked the presence of several trypanosomatids species, being *Lotmaria passim* and *Crithidia mellificae* the most prevalent ones, with this increase in mortality. Although previous studies have shown that trypanosomatid infection reduces the lifespan of bees, there is little information regarding their development in the gut when honeybees become infected. Here, for the first time we describe the haptomonad morphotype of these two trypanosomatid species in *A. mellifera*. The most characteristic feature of haptomonads is the extensive remodelling of the flagellum and the formation of junctional complexes at the host gut wall. The presence of this morphotype in the honeybee hindgut increases our understanding of the life cycle of these species and their possible pathogenic mechanisms. We found that they can multiply while attached and that their disposition, covering the hindgut walls, could hinder host nutrient uptake and consequently, represent a pathogenic mechanism itself. This attachment could also be a key stage in the life-cycle to prevent the trypanosomatids leaving the host prematurely, ensuring transmission through infective morphotypes.

## Introduction

Honeybee health is a major ecological, agricultural and societal concern due to the critical role of these insects in maintaining the balance of ecosystems and in plant reproduction. Among the wide range of abiotic and biotic stressors that affect *Apis mellifera*, parasites are one of the factors that have driven bee losses over the past decade [1]. Trypanosomatids (Euglenozoa, Kinetoplastea, Metakinetoplastina, Trypanosomatida) are one of the most extensive and successful groups of parasites in nature [2]. These organisms could compromise the fitness of their invertebrate hosts, affecting their behaviour and survival, particularly under stressful environmental conditions [2]. Since trypanosomatids were first detected in honeybees[3–5] and the first species infecting *A. mellifera* were described [6,7], their true effects on honeybee health have yet to be fully defined. However, the interest in trypanosomatids has increased greatly of late due to their possible role in bee mortality [8–10], their high prevalence [11–13] and their possible implication in honeybee colony losses[8,9,14–16].

*Crithidia mellificae* and *Lotmaria passim* are the most prevalent species and the ones that have been typically described in honeybees, although it was recently reported that other pollinator genera could also host these species, such as *Bombus* and *Osmia* [17,18]. In addition, different species of trypanosomatids have been recently detected in *A. mellifera*, some of them more common to other hosts, such as *Crithidia bombi* [19], *Crithidia expoeki* or *Crithidia acanthocephali* [13]. Hence, these organisms seem to be able to infect several host species due to their low host-specificity. The analysis of the gut from laboratory-reared honeybees showed that stress conditions (poor nutrition, an asocial context and lack of environmental exposure to microbiota) could make them more susceptible to infection by *L. passim*, while also causing alterations in behaviour and development [20]. However, not much is known about the life cycles, the pathogenic mechanisms or the transmission of trypanosomatids between different hosts.

Trypanosomatid parasites are characterized by common morphological traits, such as the presence of a kinetoplast and a single flagellum that are involved in cell signalling, motility and morphogenesis [21–24]. Stage-specific cell morphogenesis and extensive flagellar remodelling have been associated with parasite differentiation, survival and transmission [25–27]. These processes have been described previously in insect-specific haptomonad stage of *Paratrypanosoma confusum* [28] and *Crithidia fasciculata* [29] in mosquitoes, or that of *Leishmania* species in the stomodeal valve of phlebotomine sandflies [30,31]. The haptomonad stage has been described as non-motile, dividing, oval-shaped cells that are highly adhesive due to flagellar shortening and the formation of an adherent structure called the “attachment plaque” [28–30]. However, ultrastructural information of these adhesion plaques and how these species progress towards the haptomonad stage differentiation at the parasite/host interface have been poorly described for honeybee trypanosomatids.

Previous *in vivo* studies on *C. mellificae* and *L. passim* described “spheroid” and “flagellated” forms in the hindgut of honeybees following experimental infection [7,32], but no other morphotype has been described to date. Here, we describe the haptomonad cells in the hindgut of *A. mellifera* following experimental infection with *C. mellificae* choanomastigotes and *L. passim* promastigotes, adding to our understanding of the life cycle of these species. Furthermore, the presence of hemidesmosome-like binding complexes in these two species suggests that this haptomonad stage is a common trait used by trypanosomatid species when colonizing the digestive tract of honeybees.

## Methods

### Cell cultures

Reference trypanosomatid strains of *C. mellificae* (ATCC 30254) and *L. passim* (ATCC PRA403) were used for the inoculums. Cells were cultured in Brain Heart Infusion Broth (BHI: Sigma) supplemented with 10% of Heat Inactivated Fetal Bovine Serum (FBS: Gibco) and 1% of Penicillin/Streptomycin (17-602E, Lonza), as described previously [32,33]. Cultures were maintained at 27 °C in 25 cm^2^ flasks (Corning), with an initial concentration of 10^5^ cells/mL. Serial cultures were established to infect the bees with both species at the same stage of development.

### Experimental infection of honeybees

Brood frames from controlled honey bee colonies were kept at 34 ± 1 °C, and new born worker bees were caged after their emergence (10 bees/cage, 3 cages per treatment) and maintained for two days at 27 °C [32,34]. Inoculums were obtained from serial cell cultures, choanomastigote forms for *C. mellificae* at 96-168h and *L. passim* promastigote forms at 144-192 h, these times corresponding to the same stage of development (late logarithmic phase according to growth curves: unpublished data). The cells were counted in a Neubauer chamber and their concentration was adjusted to 5 x 10^4^ cells/µL with phosphate buffered saline (PBS). Two-day-old bees were starved for 2 hours and inoculated individually, as described previously [32,34,35], with either 2 µL of *C. mellificae* or *L. passim* inoculum per bee, while control bees received the same volume of PBS. Transmission Electron Microscopy (TEM) sample preparation could lead to the loss of trypanosomatids and so, to ensure accurate images were obtained from bees, they were inoculated twice a day over 12 consecutive days and after the final dose, they were fed *ad libitum* with 50% sucrose syrup + 2% Promotor L (Calier Lab). Bees infected with either species and uninfected bees were kept in separate incubators to avoid cross-contamination (Memmert® IPP500, 0.1 °C)[36].

### Gut extraction and sample preparation for microscopy

After a 12 days infection, bees were sedated with CO_2_ to extract their digestive tract by pulling the stinger [32,34,35]. Once extracted, the guts were washed with PBS to be placed and keep them stretched in 0.45 µm cellulose nitrate filters (Sartorius Biolab). Half of the bees in each cage and group were studied by light microscopy while the rest were processed for TEM. Samples for light microscopy were fixed in 10% buffered formalin for 24 hours and then embedded in paraffin, staining the microtome sections obtained (4 µm: Leica® 2155) with hematoxylin-eosin [36,37].

For TEM, the guts on the cellulose filters were fixed for 45 minutes in Karnovsky fixative (4% paraformaldehyde, 2.5% glutaraldehyde, Sorensen buffer) at 4 °C, washed and maintained in PBS until they were processed. Samples were stained with 1% osmium tetroxide (1 h), washed (x3) in distilled water, dehydrated in graded acetone series (30% - 100%, 10 min each) and embedded in graded SPURR resin-acetone series. After leaving the tissue overnight in pure SPURR resin, the ileum and rectum were separated and placed in resin blocks for ultramicrotome processing. Semithin sections (0.5 µm) were obtained (Reichert-Jung Ultracut E microtome, Leica, Wetzlar, Germany), stained with methylene blue (1%) in sodium borate water (4%) and observed under an Olympus Vanox photomicroscope (Olympus Optical Co. Ltd., Tokyo, Japan). The areas of interest were selected and trimmed, obtaining ultra-thin sections (60 nm). Grids were dual-contrasted with uranyl acetate (2%) in water and lead citrate Reynolds solution (10 min each), and then analysed and photographed on a Jeol 1010 Electron Microscope at an accelerating voltage of 80-100 kV. Cell cultures used for inoculation were also fixed for TEM analysis as described for the bee samples. For Scanning Electron Microscopy (SEM) analysis, the trypanosomatids were left for 4 hours on 13 mm round coverslips (Marienfeld) and then fixed for 24 hours at 4 °C with glutaraldehyde (2.5%) in cacodylate buffer containing 0.1 M sucrose. Subsequently, the samples were dehydrated in graded ethanol series, desiccated in a critical point dryer (Leica EM CPD 300) and evaporated with a high vacuum carbon coater (Emitech K975X). Finally, were carbon-coated for 3 minutes and observed with a ZEISS Supra microscope.

## Results

### Ultrastructural analysis of the *in vitro*-cultured flagellated forms

The ultrastructural and morphological differences between *C. mellificae* and *L. passim* species were defined through a SEM and TEM analysis of the cells *in vitro*. As previously mentioned, *C. mellificae* showed a typical choanomastigote morphology, with narrow lateral grooves at the surface and a rounded posterior end (Fig 1A). A long, free, single flagellum is inserted at the anterior apical end of the cell into a narrow flagellar pocket, extended approximately within half of the length of the cell body (Fig 1B, C). The flagellum exhibits a typical 9 x 2 + 2 axonemal pattern, with 9 pairs of peripheral microtubules surrounding 2 central microtubules (Fig 1B), that came from the flagellar body, before the kinetoplast. As indicated, the flagellum is surrounded by the reservoir or flagellar pocket (Fig 1B-D), in which some vesicles could be observed (Fig 1B, C). The kinetoplast was located just beyond the origin of the flagellum, slightly anterior or parallel to the nucleus (not shown), and enclosed by a membrane that continues with the mitochondrial network (Fig 1C). The nucleus lies in the middle-third of the cell, with visible accumulations of chromatin (Fig 1B, D). Some other structures could also be observed (Fig 1B-D), such as glycosomes, lipid droplets, acidocalcisomes, ribosomes and rough endoplasmic reticulum (RER).

**Fig 1.**
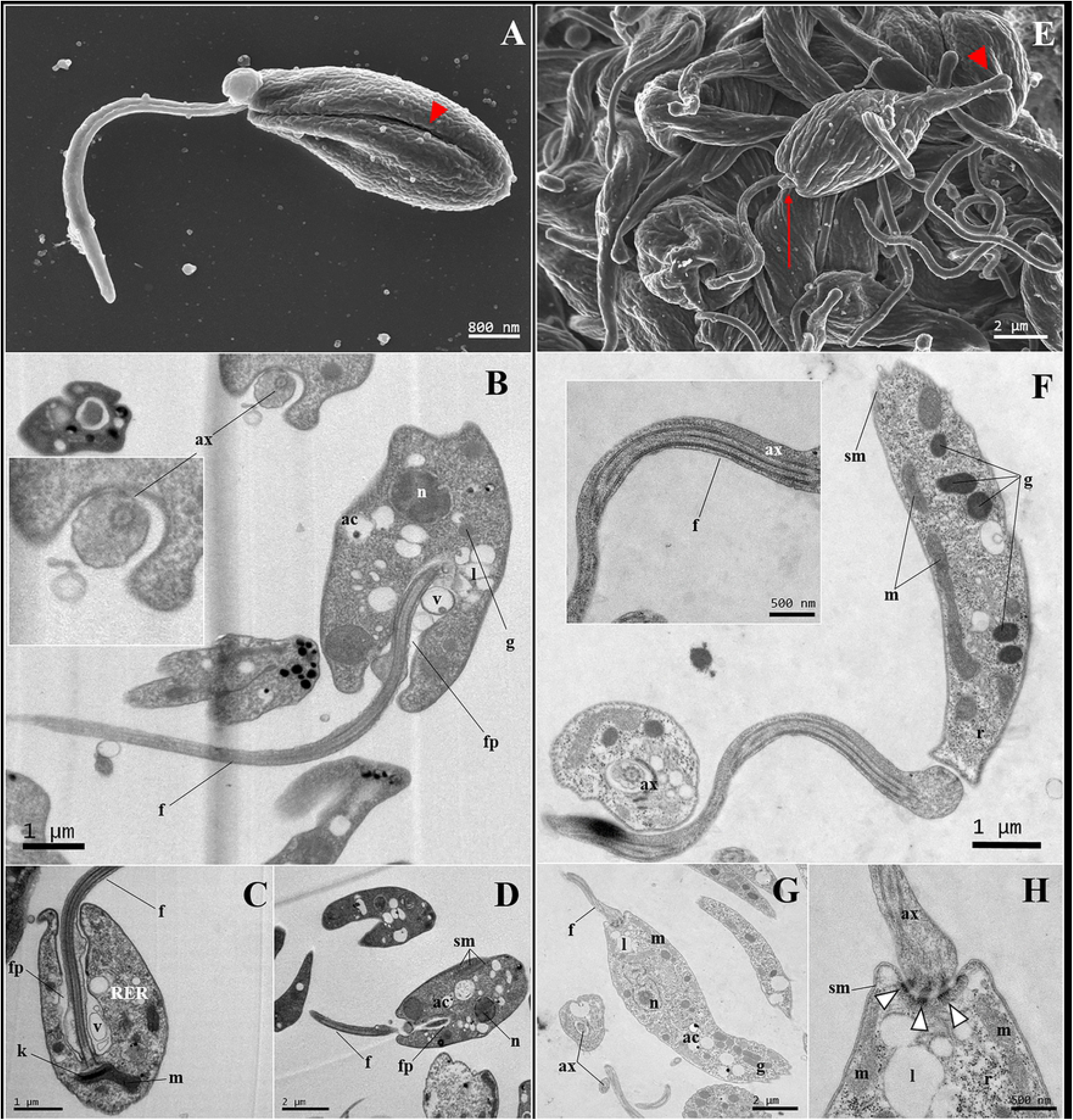
SEM (A, E) and TEM (B-D, F-H) images of cell cultures of *C. mellificae* (A-D) and *L. passim* (E-H). (**A)** Typical choanomastigote of *C. mellificae*, with an anterior insertion of the flagellum and narrow grooves across the cell body (red arrowhead). (**B-D)** Longitudinal sections of choanomastigotes in which the cell shape and organelles can be seen. (**E**) Promastigotes of cultured *L. passim* cells forming a cluster. The characteristic “nose” of the posterior end can be seen (red arrowhead), as well as the spicule at the flagellum insertion point (red arrow). (**F-H**) Longitudinal sections of promastigotes with the single, free and long flagellum, and its point of union with the cell body forming desmosome-like junction complexes (type A desmosomes: white and black arrowheads). Abbreviations: acidocalcisome (**ac**), axoneme (**ax**), flagellum (**f**), flagellar pocket (**fp**), glycosome (**g**), kinetoplast (**k**), lipids (**l**), mitochondrion (**m**), nucleus (**n**), ribosomes, (**r**), rough endoplasmic reticulum (**RER**), subpellicular microtubules (**sm**), vesicles (**v**).

By contrast, *L. passim* cells have a more lanceolated morphology, with smooth, wide and deep lateral grooves along the entire cell length (Fig 1E). A marked extension of the posterior area could also be seen, resembling a “nose” (Fig 1E). The single flagellum, of at least the length of the cell body (Fig 1F), arises from the anterior portion adopting the same axoneme structure as in the *C. mellificae* choanomastigotes (Fig 1F). In some figures, a spicule can be seen at the insertion point (Fig 1E). The flagellum is surrounded by a flagellar sheath that forms a pocket, and the attachment of both is through desmosome-like structures known as type A desmosomes (Fig 1G, H) [38]. The organelles identified in the *C. mellificae* cells were also be observed in the *L. passim* promastigote cells (Fig 1F-H).

### Microscopy study of the honeybee hindgut samples

#### Light Microscopy

Following a 12 day inoculation both species colonized the honeybee hindgut (ileum and rectum), whereas no trypanosomatids were observed in uninfected control bees. *C. mellificae* apparently showed greater tropism for the ileum (Fig 2A, B) although they were also found in the rectum (Fig 2C). By contrast, *L. passim* more frequently colonized the rectum (Fig 2D-F) although it too was also observed in ileum (Fig 2D). Both species appeared to adopt a different morphology when colonizing the ileum or rectum (Fig 2A, C-D). Trypanosomatids covered the host-epithelial surfaces, forming a carpet-like layer (Fig 2A, C, F), as well as sometimes forming clusters (Fig 2B). There were also free forms in the lumen (Fig 2A, D-E), of which some orientated the flagella towards the host surface (Fig. 2B, E). Apparently, *C. mellificae* showed multilayer clusters or carpets covering the host cells (Fig 2B), while *L. passim* seemed almost always to be arranged in a single cell layer (Fig 2F). In no case the pictures suggested that significant histological changes of the host’s epithelial cells were produced by these trypanosomatids.

**Fig 2.**
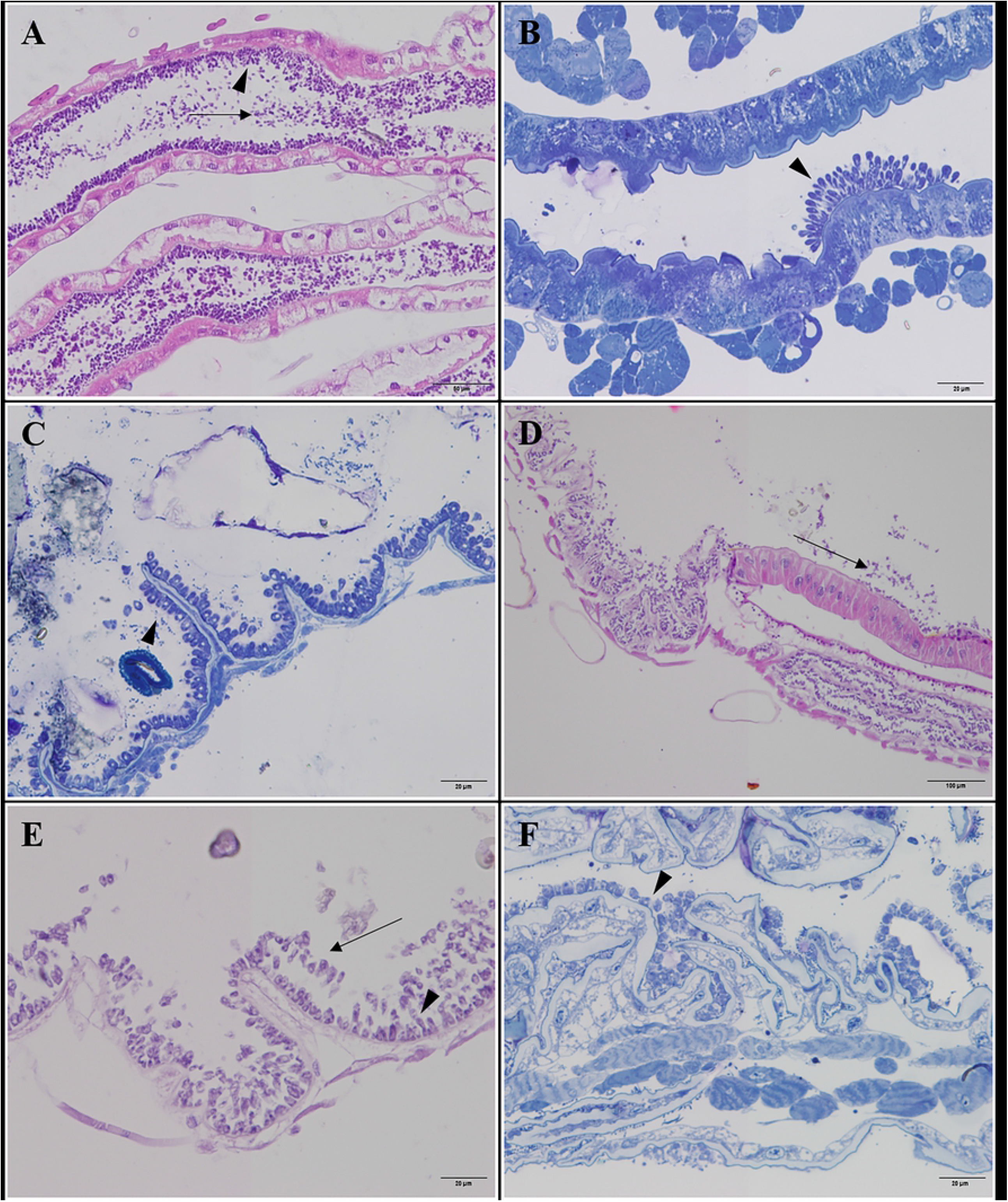
Light microscopy images from bees infected with *C. mellificae* (A-C) and *L. passim* (D-F). **(A)** Haematoxylin-eosin (H&E) stained longitudinal section (Magnification: 20X) of the ileum from a *C. mellificae* infected bee, in which free forms and attached forms can be seen. (**B**) Methylene blue-stained longitudinal semi-thin section (40X) from the ileum of a bee infected with *C. mellificae* cells that can be seen to form a cluster, and (**C**) a rectal longitudinal methylene blue-stained semi-thin section (40X) from a *C. mellificae* infected bee in which the trypanosomatids appear to form a single layer of cells covering the rectal surface. (**D**) H&E stained longitudinal section (10X) of the connection of the ileum and rectum of a *L. passim* infected bee in which two different morphotypes could be seen, ones covering the gut surface and others free in the lumen. (**E**) Longitudinal H&E stained section (40X) of the rectum of a bee infected with *L. passim* promastigotes. (**F**) Methylene blue-stained semi-thin longitudinal section (40X) of the rectum of a *L. passim* infected bee. The cells form a “carpet” covering the host epithelial cells. Leyend: (**arrowhead**) attached trypanosomatids covering the hindgut walls, (**arrow**) free flagellated forms in the lumen.

#### Transmission Electron Microscopy (TEM)

TEM images of *C. mellificae* were obtained mostly from the ileum (Fig 3A-E), and only in one of the samples could be seen in the rectum (Fig 3F-H), whilst *L. passim* was only observed in the rectum (Fig 4) no TEM images from ileum were obtained for this species.Both trypanosomatid species attached to the host epithelial surface (Fig 3A and Fig 4A), although the cell morphology differed to that seen in the cultures used for inoculation (choanomastigotes and promastigotes). In both cases, the cells presented a haptomonad morphotype (Fig 5), which have a round-to-bell shape, and a remarkable remodelling of the flagellum, that became an attachment pad. This allows them to remain attached and thrive in the host hindgut (Fig 3B, D, H and Fig 4B, C, F). The high density of cells in close contact lining the epithelium appears to make them adapt their shape to the space available (Fig 3A, D, E and Fig 4A, C, F). In both species, parasites maintained close contact with either the host epithelia or with the cell clusters, which suggest host-parasite and parasite-parasite interactions. In accordance with the light microscopy analysis, none of the TEM images provided evidence that the trypanosomatids produced host cell damage. As flagellum remodelling was the most apparent phenomenon it will be analysed separately.

**Fig 3.**
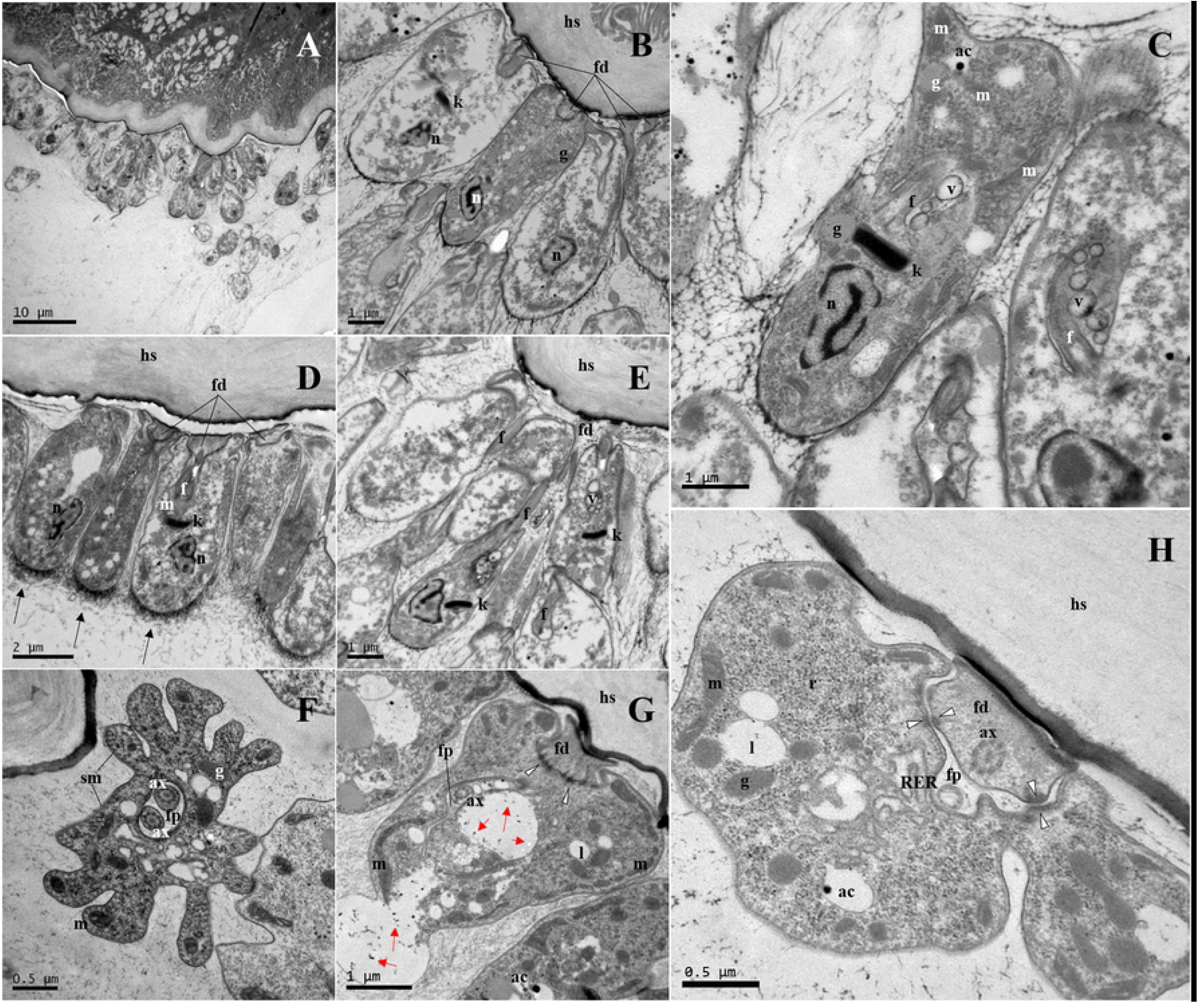
TEM images obtained from the ileum (A-E) and rectum (F-H) of *C. mellificae* infected bees. (**A**) Longitudinal section of one of the clusters found in the ileum where some free trypanosomatid forms could be seen. (**B-D**) Longitudinal sections of the haptomonad adherent forms found on the ileum walls, with the modified flagellum that allowed the attachment. The kinetoplast is situated anterior to the nucleus. Outside the trypanosomatids, electron-dense material could be seen, especially in the posterior part of the cells (black arrows). (**E**) Longitudinal section of the different cell morphotypes, some of them unattached cells that had a typical unmodified flagellum, although it seemed to be oriented towards the host surface which indicates tropism towards it. (**F)** Cross-section of one of these haptomonads from the rectum in which two flagellar axonemes could be observed, indicating a process of cell division. (**G, H)** Longitudinal section of *C. mellificae* haptomonad forms. Some of the cells (**G**) seem to be expelling material of different electron-density (red arrows) inside vesicles. Junction complexes could be observed between the flagellapodia and the cell body (type A desmosomes: black and white arrowhead). Abbreviations: acidocalcisome (**ac**), axoneme (**ax**), flagellum (**f**), flagellopodium (**fd**), flagellar pocket (**fp**), glycosome (**g**), host surface (**hs**), kinetoplast (**k**), lipids (**l**), mitochondrion (**m**), nucleus (**n**), rough endoplasmic reticulum (**RER**), subpellicular microtubules (**sm**), vesicles (**v**).

**Fig 4.**
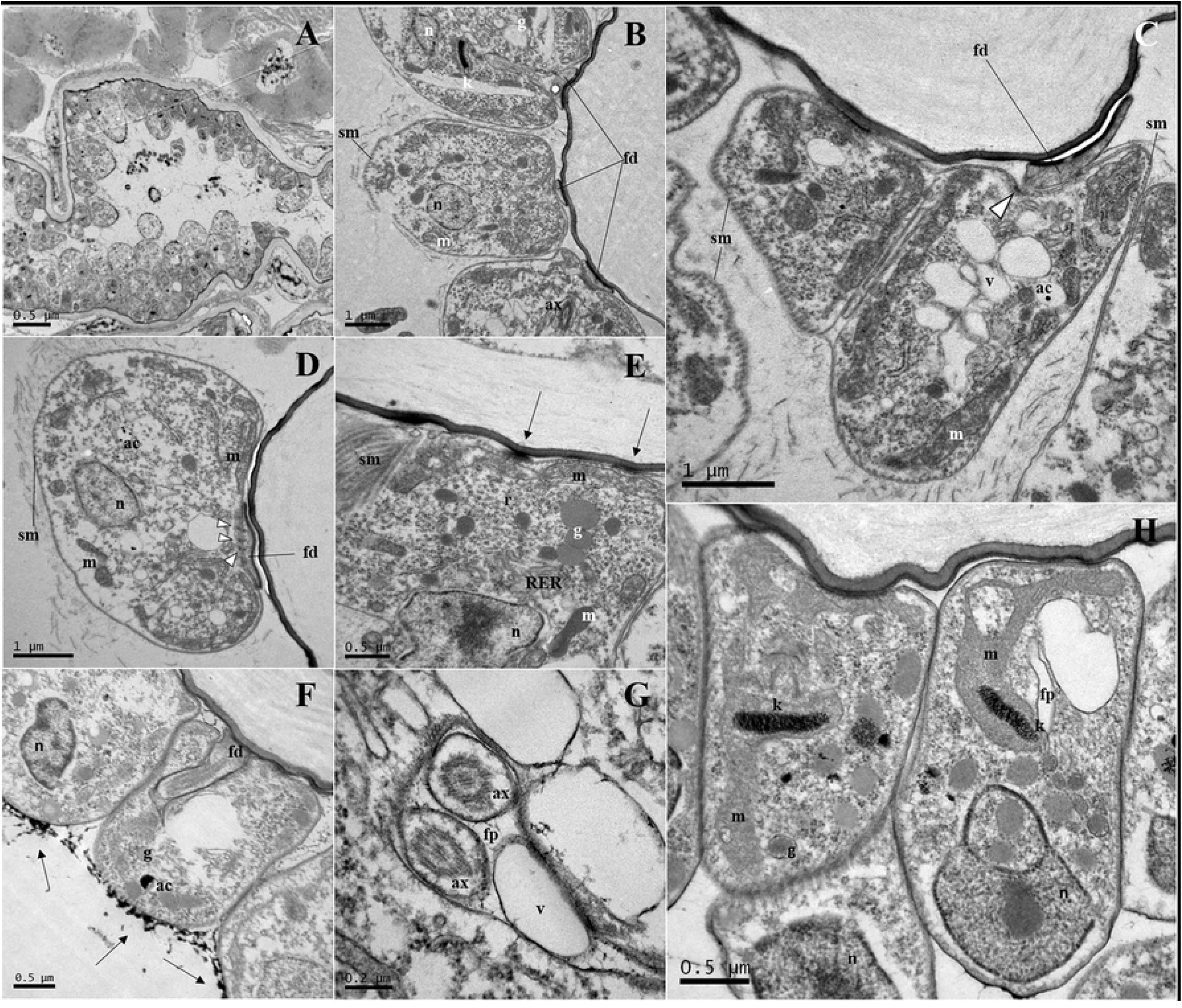
TEM images of the rectum of *L. passim* infected bees. (**A)** Longitudinal section of the rectum wall where the disposition of the cells in the lumen is shown. Some of the cells seem not to be attached to the surface of the host epithelium. (**B-D, F, H**) Longitudinal sections of the haptomonad forms found in the rectum. The flagellapodia could be seen in almost all the images, showing their adaptability to the host surface, and the desmosome-like complexes that join this structure to the cell body are also evident (black and white arrowheads). The expelled material of unknown nature is seen outside these cells and accumulated in the posterior part of the cells (black arrows). The relative position of the nucleus and the kinetoplast is also shown in some images (**B, H**), and many of the organelles typical of trypanosomatids were seen. (**E)** Detail of a longitudinal section of the cell membrane contact zone and not that of the flagellopodium membrane with the host surface, where electron-dense points could be seen. The single layer of subpellicular microtubules is situated beneath the membrane. (**G**) Cross-section of the haptomonad form showing two flagella entering into the flagellar pocket, reflecting an early stage of cell division. Abbreviations: acidocalcisome (**ac**), axoneme (**ax**), flagellopodium (**fd**), flagellar pocket (**fp**), glycosome (**g**), kinetoplast (**k**), mitochondrion **(m)**, nucleus (**n**), rough endoplasmic reticulum (**RER**), subpellicular microtubules (**sm**) vesicles (**v**).

**Fig 5.**
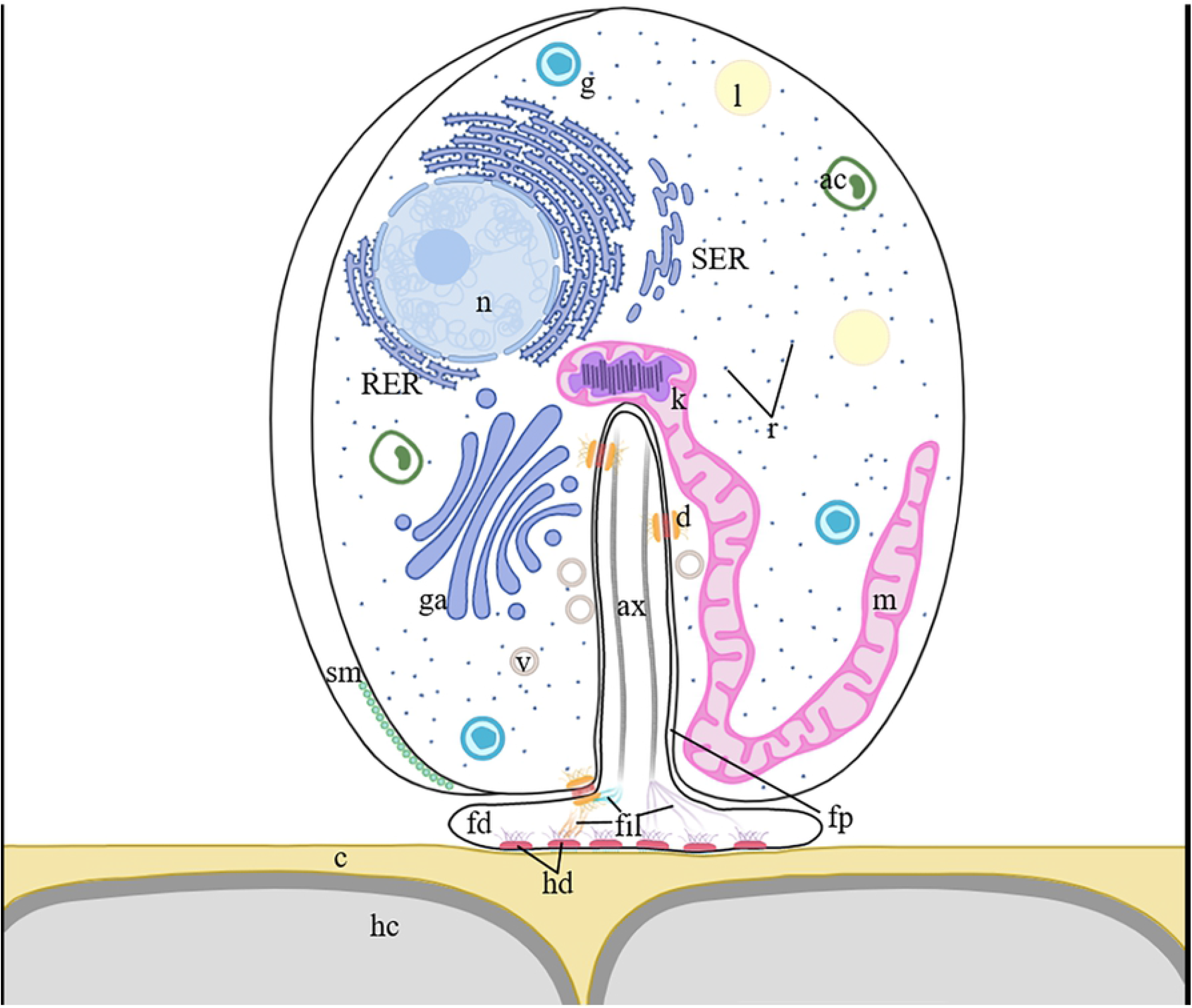
Scheme representing the haptomonad morphotype found in the hindgut of bees infected with *C. mellificae* and *L. passim*. Abbreviations: acidocalcisome (**ac**), axoneme (**ax**), cuticle (**c**), desmosome (**d**), filaments (**f**), flagellopodium (**fd**), flagellar pocket (**fp**), glycosome (**g**), golgi apparatus (**ga**), host cells (**hc**), hemidesmosome-like complex (**hd**), kinetoplast (**k**), lipids (**l**), mitochondrion (**m**), nucleus (**n**), ribosome (**r**), rough endoplasmic reticulum (**RER**), smooth endoplasmic reticulum (**SER**), subpellicular microtubules (**sm**), vesicles (**v**).

In general, the haptomonad nucleus of both species was oval or slightly elongated, middle-third located towards the aflagellar pole of the cell, with heterochromatin accumulated in the membrane (Fig 3B-D, E and Fig 4E, F). The kinetoplast was situated anterior or laterally to the nucleus (Fig 3C-E and Fig 4B, H), and its connection with the unique, large, tubular and peripheral mitochondrion could be seen in some cases (Fig 4H). The flagellar pocket starts immediately anterior to the kinetoplast and similarly to what was observed in the *in vitro* morphotypes, the flagellar pocket seems to be inserted approximately into half the length of the cell body in *C. mellificae* (Fig 3C, D). By contrast, *L. passim* had a smaller flagellar pocket located towards the anterior end of the cell body (Fig 4B). The flagellum originated inside the pocket, emerging at the anterior part of the cell as a “flagellopodium” or “attachment pad”, and forming an electron-dense junction with the host’s cell surface (Fig 3B, C, G, H and Fig 4B-D, F, H). However, contact points of the flagellopodium surface with the host epithelium were observed as electron-dense areas in some images, yet there were no signs of the formation of junction complexes (Fig 4E).

Other cell structures observed included large numbers of ribosomes distributed throughout the cytoplasm, RER, as well as other organelles with different electron densities like acidocalcisomes, glycosomes, lipid droplets and vesicles (Fig 3C, H, G and Fig 4B, D, E). Moreover, we observed a single layer of subpellicular microtubules (Fig 3H, F and Fig 4B, D, E). Some cells contained electron-lucid secretion, with visible electron-dense particles of uncharacterized nature, that appeared to be released through the posterior membrane (Fig 3G). In fact, these electron-dense particles could also be observed in the lumen, in particular in the extracellular space around the posterior part of both *C. mellificae* and *L. passim* cells (Fig 3D and Fig 4F). Different events of the division cycle of the cells were seen, with some of them with more than one flagella (Fig 3F and Fig 4G). Despite the layout of the cells, in close contact with each other, no plasma membrane fusion appeared to occur and the cells maintained their own membrane intact (Fig 4H). Non-attached trypanosomatids could also be observed and as in the light microscopy images, most of them were oriented with the flagella towards the epithelial surface (Fig 3E and Fig 4A).

The attachment of trypanosomatids to the honeybee hindgut is mediated by a flagellar structure that here, according to previous works [39] is called “flagellopodium” (sing.) or “flagellapodia” (pl.), and was regularly found in all the images (Fig 6 and Fig 7). Both *C. mellificae* and *L. passim* haptomonad forms showed a similar flagellopodium, with a typical 9 x 2 + 2 microtubules conformation (Fig 6A, D, E and Fig 7A, B, D). The extension and length of the flagellopodium varied according to the location of the trypanosomatid to the host (Fig 6B and Fig 7C, D). While *C. mellificae* appeared to have a longer flagellopodium (Fig 6A-B), *L. passim* seemed to flatten it to extend it and cover a larger host cell surface (Fig 7B-C) although no quantitation was performed.

**Fig 6.**
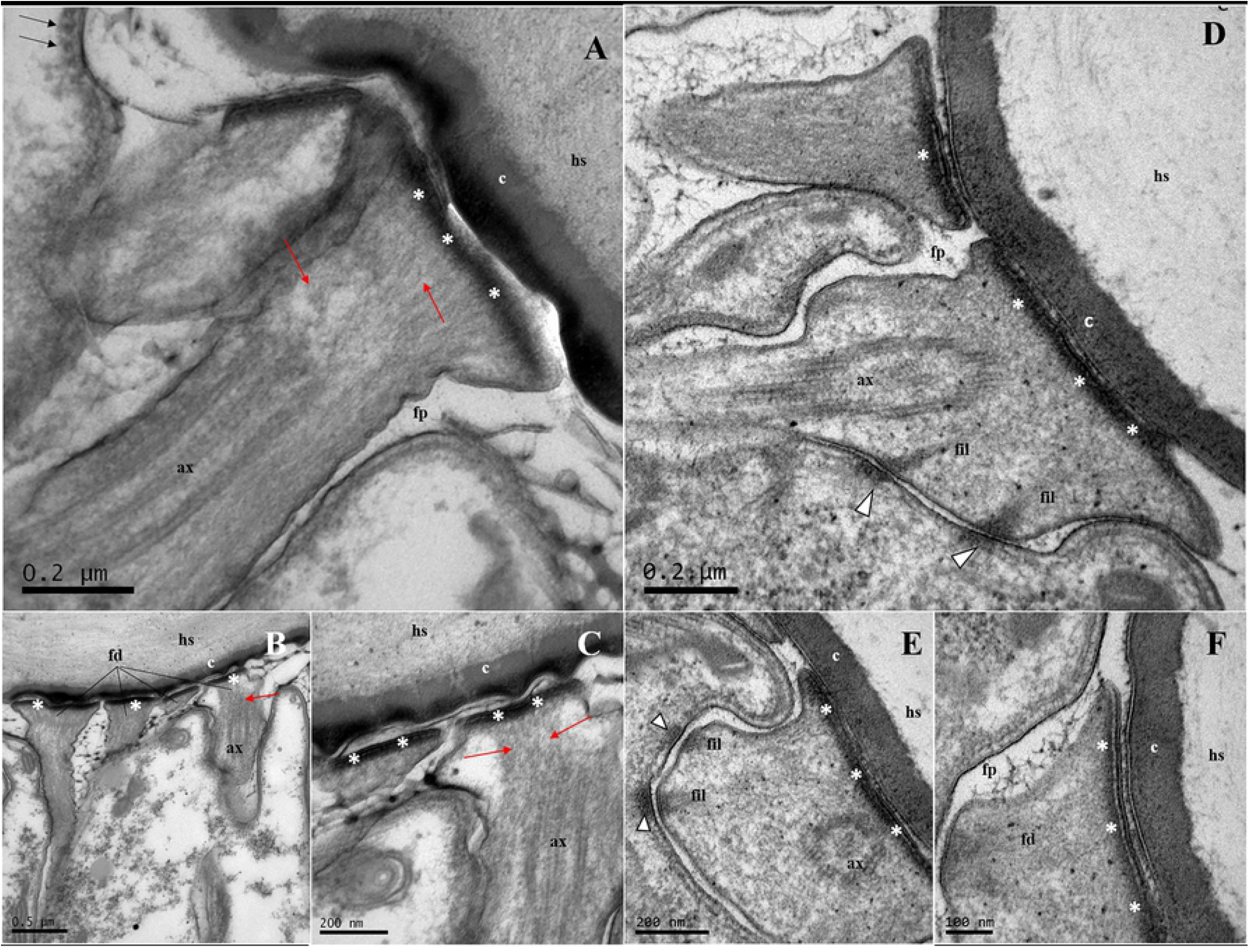
Detail of the flagellapodia of the *C. mellificae* haptomonad form found in the ileum (A-C) and rectum (D-F). **(A)** The single layer of subpellicular microtubules on the cell surface is shown (black arrows). Two types of junction complexes could be seen (**D, E**): the desmosome-like junction complexes (type A desmosomes) between the flagellapodia and the flagellar pocket membrane (black and white arrowheads); and the hemidesmosome-like junctions between the bulge of the flagellapodia and the host surface, with electron-dense material beneath the inner leaflet (asterisks). Longitudinal sections of flagellapodia showed the mass of filaments that reinforce the complex, connecting both kinds of junctions (**fil**), and flagellar cytoskeleton filaments can be seen connecting the axoneme with the hemidesmosome-like complex (red arrows). Flagellapodia of different cells in close contact with each other that help attach them to the host surface could also be observed. **(B)** Flagellapodia attached to the rectum (**D-F**) seem to present a different morphology, taller and thinner than those in the ileum that were wider and more extended. (**F)** Higher magnification of the *C. mellificae* host surface and the flagellopodium contact point. Abbreviations: axoneme (**ax**), cuticle (**c**), filaments (**fil**), flagellopodium (**fd**), flagellar pocket (**fp**), host surface (**hs**).

**Fig 7.**
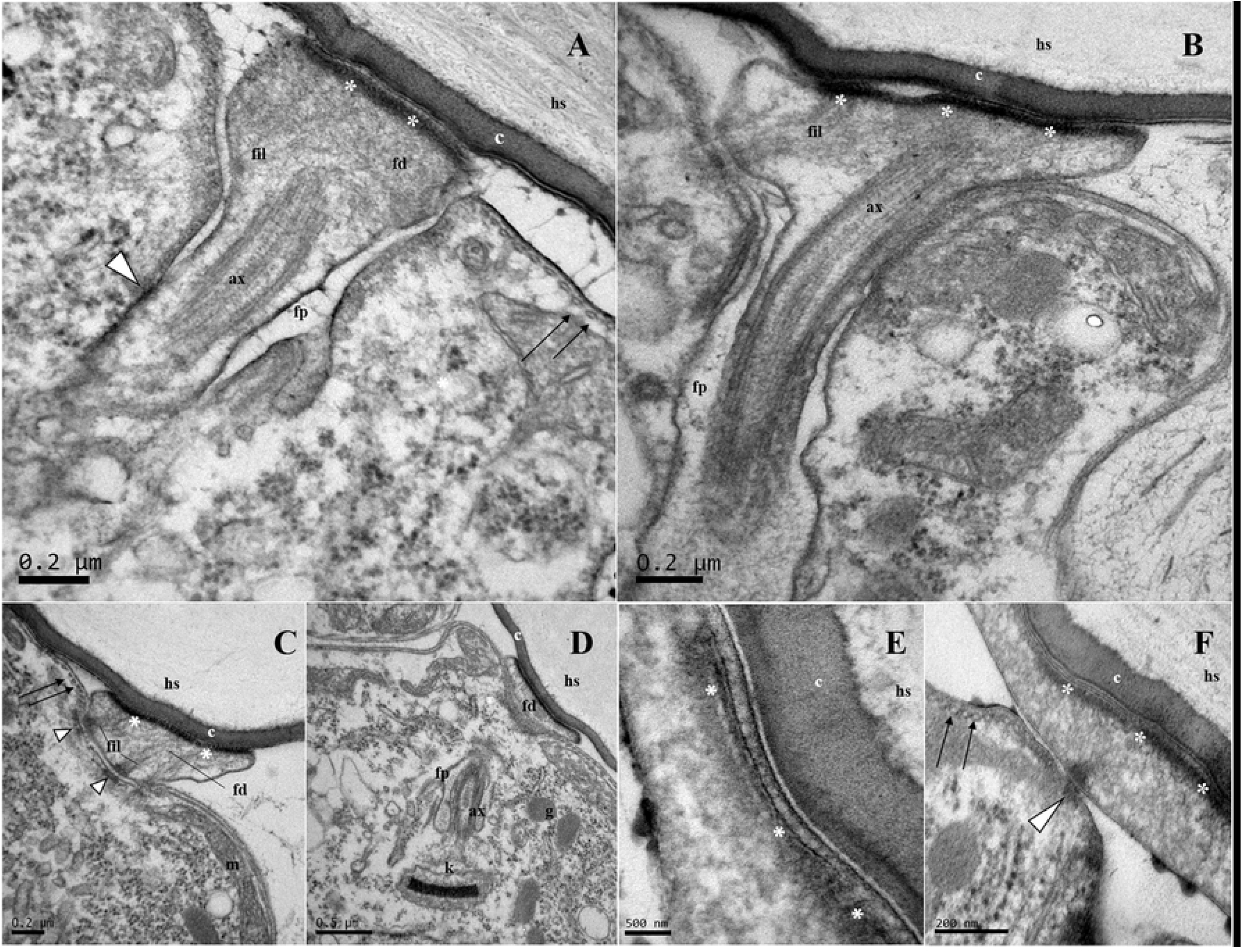
Longitudinal section of the flagellapodia of the haptomonad forms from the rectum of *L. passim* infected bees. In these sections the two types of junction complexes could be observed: type A desmosomes (black and white arrowheads) and the electron-dense material of the hemidesmosome-like structures (asterisks). The filament network connected to the flagellar structure could also be seen (**B, C**). Other organelles and structures were evident, such as the kinetoplast, the start of the flagellar apparatus inside the cell (**D**) and the subpellicular array of microtubules (black arrows) (**A, C, F**). Abbreviations: axoneme (**ax**), cuticle (**c**), filaments (**fil**), flagellopodium (**fd**), flagellar pocket (**fp**), glycosome (**g**), host surface (**hs**), kinetoplast (**k**).

The modified flagellum contacts the intestinal cells through hemidesmosome-like structures seen as electron-dense areas in the images (Fig 6D, E and Fig 7B, D, F). The main flagellar remodelling occurred at the distal region, where the haptomonad forms an attachment pad to join the epithelial cells via large bulges at the base of the flagellopodium (Fig 6C and Fig 7B). The surface of this structure was aligned with and extended over the host cells, creating a point of contact between the trypanosomatid and host that was mediated by the formation of hemidesmosome-like junction complexes. These complexes had a higher electron-density beneath the flagellar membrane (Fig 6A, D, F and Fig 7B, E, F). These junctions between the flagellapodia and the honeybee hindgut surface lay at a critical distance of 10–20 nm, and the gap between the two membranes was filled by a well-packed electron-dense array of filaments (Fig 6A, E, F and Fig 7E), forming a strip-like zonular hemidesmosome. In the images of both species, no tonofibril-like arrays were evident at the surface or at a distance, highlighting the apparent absence of cytoskeletal connections on the host side. However, an array of filaments reinforcing the entire flagellopodium complex was evident, connecting the hemidesmosome-like junctions, the type A desmosomes and the axoneme (Fig 6 and Fig 7A-C).

## Discussion

The present work describes a new trypanosomatid morphotype in honeybees, the haptomonad form, not previously described for *L. passim* and *C. mellificae*, the most prevalent species infecting honeybees. This haptomonad morphotype involves flagellar remodelling that allows these organisms to remain attached to the hindgut walls.

The term “haptomonad” was first used to name the “gregariniform phase” attached to the *Culex pipiens* stomach [40] and then later to refer to the attached stage of development of several trypanosomatid species in different hosts [41–43]. The haptomonad morphotype is indeed a regular pattern of the trypanosomatid life cycle in the digestive tract of insects [28,41,44], and it can also be seen *in vitro* attached to plastic surfaces and other materials [45,46]. Spheroid non-flagellated trypanosomatids have previously been described attached to the rectum and ileum walls of honeybees [7,32], but to the best of our knowledge this is the first time that haptomonad forms have been seen in honeybees. Moreover, both species studied showed the haptomonad morphotype irrespective of the hindgut region analyzed. However, we should emphasize that these morphotypes were observed under our experimental conditions, for which the inoculum characteristics, as well as the number of culture passages, the morphotype used or the age of the cell cultures should be borne in mind [32,33].

The most remarkable feature of these haptomonad forms is the flagellar remodelling that results in the formation of the flagellopodium, strikingly different to the *in vitro*-cultured flagellated forms of the inoculum. As a consequence of this flagellar remodelling, the entire cell has to adapt its morphology [24]. It is also noteworthy that two main junction-complexes could be seen in almost all the images. First, Type A desmosomes form between the flagellopodium sheath and the flagellar pocket that connect the modified flagellum to the trypanosomatid cell body, a structure that can also be seen in the free-flagellated forms. The number of these unions seems not to be constant, in accordance with earlier studies [44]. The other remarkable junction complex is the hemidesmosome-like junction between flagellapodia and the host gut surface. Previous studies suggested that hemidesmosomes are the optimal junctions to attach to smooth surfaces like the cuticle [47]. In the images is shown as an electron-dense zone that extends along the membrane of the flagellar bulge is evident, which is in close contact with the host cell. However, according to earlier studies [41] there would not be one large junction zone but a hemidesmosome at each point of contact, and these attachment points will be determined by the distance between the host surface and the flagellopodium membrane [38,41]. Unfortunately, the magnification of the images obtained does not allow us to differentiate this, so further studies would be of interest to address this issue. In other trypanosomatid species, such as *Leishmania mexicana*, the formation of hemidesmosome junctions is preceded by the expansion of the flagellar sheath, a process that could break the membranes, releasing phospholipids that could form the hemidesmosome mass [41]. Nevertheless, to the best of our knowledge the nature of this adhesion remains unclear and as such, it could be chemical or electrical due to the charge difference between the membranes [48]. In addition, bands of filaments were seen between the hemidesmosome-like structures and the type A desmosomes of the flagellar pocket [25,45], connecting these two kinds of junctions and thereby strengthening the entire flagellum complex, consequently securing the attachment.

It is known that the life cycle of trypanosomatids encompasses controlled and adaptive plasticity to change their cell morphology, as well as their biochemical properties, in order to thrive and respond to changing environmental conditions [49,50]. In our experimental conditions there are two possible sites of colonization, the ileum and rectum, since only a few free-flagellated trypanosomatid cells were seen in the midgut and none of these were attached. By contrast, haptomonad cells were found covering the honeybees’ hindgut surfaces, with flagellated forms around and among them, as seen previously [7]. Each hindgut region has is specialized for different functions and consequently, they have different conditions, including variations in their microbiota and gut cell walls [51–53]. The ileum acts as a connector between the midgut and the rectum, and although its function is not entirely clear, the composition of its epithelium suggests that it may be involved in water and nutrient absorption. By contrast, the rectum is mainly responsible for the absorption of water and minerals, regulating osmotic pressure. It also releases some enzymes, such as catalase, to degrade the toxic H_2_O_2_ produced by glycolic acid synthesis, and it stores faecal waste until defecation [54–56]. Although we observed both species in the two regions of the hindgut by light microscopy, the differences between these gut regions could be a key factor to explain their preferential location, with *C. mellificae* tending to settle in the ileum, while *L. passim* appears to be attracted to colonize the rectum. Moreover, preceding research of *C. fasciculata* infecting *Anopheles gambiae* has suggested that the attachment and extension of the flagella, and consequently its length, is induced by cell crowding [51]. Together with the smaller extension of the ileum, this would lead to the presence of several layers of trypanosomatids attached to the gut wall and consequently, a variable length of their flagella depending on their position relative to the host [51]. This fact could explain why *C. mellificae* seems to form clusters of cells in the ileum, while *L. passim* appears to be arranged in a single layer in the rectum.

The composition and properties of the cuticular layer that coats the hindgut epithelial cells [51,57] could also cause some kind of attraction or tropism, and mediate flagellar attachment [41,44,58]. Many free, flagellated cells found in the hindgut are found with their flagella oriented towards the gut walls, presumably looking for attachment, although they could be “haptomonad daughter cells” detaching from the epithelial wall [28]. The lack of a cuticular coating, the presence of peritrophic membranes and the secretion of digestive enzymes could be factors that explain why it is not common to see attached forms in the midgut, in accordance with previous works [7,51].

The pathogenicity of insect trypanosomatids, which are often found at high densities in the gut, is not entirely clear [57]. Since both species seem to cause no pathological alterations at the host surface, it would be highly probable that they act at the lumen. Both *L. passim* and *C. mellificae* cover the hindgut walls, forming carpets, which could jeopardize metabolite absorption, as has been suggested previously [57,58], and may even block this process [51]. Alternatively, the haptomonad cells could sequester molecules and compounds, either directly from the lumen or from the intestinal cells of the bee, although further studies will be needed to clarify this. Both these processes could generate a nutritional deficit for the bee, which could increase their rates of mortality. Intriguingly, the absence of attached cells in the midgut suggests that these organisms have a tropism for posterior locations rather than the midgut [7,51], where in theory the availability of nutrients would be higher. Nevertheless, it has been proposed that the Malpighian tubes permit the passage of low molecular weight substances to the gut, like sugars and amino acids, to be absorbed by the rectum [59], providing an ideal environment for these flagellates to thrive [51]. Another potential pathogenic mechanism is that the trypanosomatids could release diverse types of molecules of diverse nature (such as proteins, metabolites or toxins) inside extracellular vesicles (EVs) [60], which might correspond to the uncharacterized material of different electron-density seen outside the cells.

This work helps shed more light on the life cycle of these two species, in particular regarding the intrahost development when a bee is infected with choanomastigotes and promastigotes, although how that cycle is completed outside the host remains unclear. As social insects, honeybee colonies are the perfect environment to establish and spread parasitic infections due to their particular traits [61], and several routes of transmission have been suggested [62–64]. Haptomonad forms are found mainly in the hindgut, yet it is unknown whether this form is that which is excreted with the faeces and thus, spreads the infection in the bee colony, or if oral-oral transmission is the most common route. In any case, attachment could be a key process to prevent non-infective morphotypes from leaving the host prematurely, as reported for other trypanosomatid species [65]. Alternatively, this attachment may serve to secure it to a substrate where the trypanosomatid could perform essential functions, such as accumulate nutrition or reproduce.

Further studies will be needed to completely understand the life cycle of these trypanosomatids and to clarify the morphotype that can survive outside the host and that can be transmitted to other bees. Likewise, precisely defining their mechanisms of pathogenicity and the biological function of adhesion through hemidesmosomes, as well as the effect of this on bee physiology, would be of great interest to better understand the relationship between these organisms.

## Acknowledgements

The authors want to thank J. Almagro, J. García, V. Albendea, C. Uceta, M. Gajero, T. Corrales, C. Botías, M. Benito, C. Jabal-Uriel and D. Aguado at the Laboratorio de Patología Apícola (Centro de Investigación Apícola y Agroambiental (CIAPA), IRIAF, Junta de Comunidades de Castilla-La Mancha), and Mr Aranda at the Departamento de Medicina y Cirugía Animal (Facultad de Veterinaria, UCM), for their technical support and help. We also thank Marisa García and Miriam González at the CNME (UCM) for their assistance in the preparation, processing and visualisation of the TEM samples. In addition, we wish to thank Instituto Regional de Investigación y Desarrollo Agroalimentario y Forestal de Castilla-La Mancha (IRIAF) for the economic support to carry out this work.

## References

1. Goulson D, Nicholls E, Botías C, Rotheray EL. Bee declines driven by combined Stress from parasites, pesticides, and lack of flowers. Science (80-). 2015;347. doi:10.1126/science.1255957

2. Maslov DA, Votýpka J, Yurchenko V, Lukeš J. Diversity and phylogeny of insect trypanosomatids: All that is hidden shall be revealed. Trends Parasitol. 2013;29: 43–52. doi:10.1016/j.pt.2012.11.001

3. Fanthan HB, Porter A. Note on certain protozoa found in bees. Suppl J Board Agricul. 1912.

4. Lotmar R. über Flagellaten und Bakterien im Dünndarm der Honigbiene (Apis mellifica). H. Sauerlander. 1946.

5. Giavarini I. Sui flagellati dell’ intestino tenue dell’ape domestica. Bolletino di Zool. 1950;17: 603–608. doi:10.1080/11250005009436845

6. Langridge DF, McGhee RB. Crithidia mellificae n. sp. an Acidophilic Trypanosomatid of the Honey Bee Apis mellifera. J Protozool. 1967;14: 485–487. doi:10.1111/j.1550-7408.1967.tb02033.x

7. Schwarz RS, Bauchan GR, Murphy CA, Ravoet J, De Graaf DC, Evans JD. Characterization of two species of trypanosomatidae from the Honey Bee Apis mellifera: Crithidia mellificae Langridge and McGhee, and Lotmaria passim n. gen., n. sp. J Eukaryot Microbiol. 2015;62: 567–583. doi:10.1111/jeu.12209

8. Ravoet J, Maharramov J, Meeus I, De Smet L, Wenseleers T, Smagghe G, et al. Comprehensive Bee Pathogen Screening in Belgium Reveals Crithidia mellificae as a New Contributory Factor to Winter Mortality. PLoS One. 2013;8. doi:10.1371/journal.pone.0072443

9. Cepero A, Ravoet J, Gómez-Moracho T, Bernal JL, Del Nozal MJ, Bartolomé C, et al. Holistic screening of collapsing honey bee colonies in Spain: A case study. BMC Res Notes. 2014;7. doi:10.1186/1756-0500-7-649

10. Stevanovic J, Schwarz RS, Vejnovic B, Evans JD, Irwin RE, Glavinic U, et al. Species-specific diagnostics of Apis mellifera trypanosomatids: A nine-year survey (2007–2015) for trypanosomatids and microsporidians in Serbian honey bees. J Invertebr Pathol. 2016;139: 6–11. doi:10.1016/j.jip.2016.07.001

11. Arismendi N, Bruna A, Zapata N, Vargas M. PCR-specific detection of recently described Lotmaria passim (Trypanosomatidae) in Chilean apiaries. J Invertebr Pathol. 2016;134: 1–5. doi:10.1016/j.jip.2015.12.008

12. Buendía, M., Martín-Hernández, R., Ornosa C, Barrios L, Bartolomé C, Higes M. Epidemiological study of honeybee pathogens in Europe: The results of Castilla-La Mancha (Spain). Spanish J Agric Res. 2018;16: e0502. doi:10.5424/sjar/2018162-11474

13. Bartolomé C, Buendía-Abad M, Benito M, Sobrino B, Amigo J, Carracedo A, et al. Longitudinal analysis on parasite diversity in honeybee colonies: new taxa, high frequency of mixed infections and seasonal patterns of variation. Sci Rep. 2020;10: 1–9. doi:10.1038/s41598-020-67183-3

14. Castelli L, Branchiccela B, Invernizzi C, Tomasco I, Basualdo M, Rodriguez M, et al. Detection of Lotmaria passim in Africanized and European honey bees from Uruguay, Argentina and Chile. J Invertebr Pathol. 2019;160: 95–97. doi:10.1016/j.jip.2018.11.004

15. Regan T, Barnett MW, Laetsch DR, Bush SJ, Wragg D, Budge GE, et al. Characterisation of the British honey bee metagenome. Nat Commun. 2018;9. doi:10.1038/s41467-018-07426-0

16. D’Alvise P, Seeburger V, Gihring K, Kieboom M, Hasselmann M. Seasonal dynamics and co-occurrence patterns of honey bee pathogens revealed by high-throughput RT-qPCR analysis. Ecol Evol. 2019;9: 10241–10252. doi:10.1002/ece3.5544

17. Graystock P, Goulson D, Hughes WOH. The relationship between managed bees and the prevalence of parasites in bumblebees. PeerJ. 2014;2: e522. doi:10.7717/peerj.522

18. Strobl V, Yañez O, Straub L, Albrecht M, Neumann P. Trypanosomatid parasites infecting managed honeybees and wild solitary bees. Int J Parasitol. 2019;49: 605–613. doi:10.1016/j.ijpara.2019.03.006

19. Bartolomé C, Buendía M, Benito M, De la Rúa P, Ornosa C, Martín-Hernández R, et al. A new multiplex PCR protocol to detect mixed trypanosomatid infections in species of Apis and Bombus. J Invertebr Pathol. 2018;154: 37–41. doi:10.1016/j.jip.2018.03.015

20. Schwarz RS, Moran NA, Evans JD. Early gut colonizers shape parasite susceptibility and microbiota composition in honey bee workers. Proc Natl Acad Sci. 2016;113: 9345–9350. doi:10.1073/pnas.1606631113

21. Langousis G, Hill KL. Motility and more: The flagellum of Trypanosoma brucei. Nat Rev Microbiol. 2014;12: 505–518. doi:10.1038/nrmicro3274

22. Vaughan S. Assembly of the flagellum and its role in cell morphogenesis in Trypanosoma brucei. Curr Opin Microbiol. 2010;13: 453–458. doi:10.1016/j.mib.2010.05.006

23. Bargul JL, Jung J, McOdimba FA, Omogo CO, Adung’a VO, Krüger T, et al. Species-Specific Adaptations of Trypanosome Morphology and Motility to the Mammalian Host. PLoS Pathog. 2016;12: 1–29. doi:10.1371/journal.ppat.1005448

24. Sunter JD, Gull K. The Flagellum Attachment Zone: ‘The Cellular Ruler’ of Trypanosome Morphology. Trends Parasitol. 2016;32: 309–324. doi:10.1016/j.pt.2015.12.010

25. Vickerman K. The Mode of Attachment of Trypanosoma vivax in the Proboscis of the Tsetse Fly Glossina fuscipes: an Ultrastructural Study of the Epimastigote Stage of the Trypanosome*. J Protozool. 1973;20: 394–404. doi:10.1111/j.1550-7408.1973.tb00909.x

26. Bonaldo MC, Souto-Padron T, De Souza W, Goldenberg S. Cell-substrate adhesion during Trypanosoma cruzi differentiation. J Cell Biol. 1988;106: 1349– 1358. doi:10.1083/jcb.106.4.1349

27. Hendry KAK, Vickerman K. The requirement for epimastigote attachment during division and metacyclogenesis in Trypanosoma congolense. Parasitol Res. 1988;74: 403–408. doi:10.1007/BF00535138

28. Skalický T, Dobáková E, Wheeler RJ, Tesařová M, Flegontov P, Jirsová D, et al. Extensive flagellar remodeling during the complex life cycle of Paratrypanosoma, an early-branching trypanosomatid. Proc Natl Acad Sci U S A. 2017;114: 11757–11762. doi:10.1073/pnas.1712311114

29. Filosa JN, Berry CT, Ruthel G, Beverley SM, Warren WC, Tomlinson C, et al. Dramatic changes in gene expression in different forms of Crithidia fasciculata reveal potential mechanisms for insect-specific adhesion in kinetoplastid parasites. PLoS Negl Trop Dis. 2019;13: 1–29. doi:10.1371/journal.pntd.0007570

30. Warburg A, Hamada GS, Schlein Y, Shire D. Scanning electron microscopy of Leishmania major in Phlebotomus papatasi. Zeitschrift für Parasitenkd Parasitol Res. 1986;72: 423–431. doi:10.1007/BF00927886

31. Sacks D, Kamhawi S. Molecular Aspects of Parasite-Vector and Vector-Host Interactions in Leishmaniasis. Annu Rev Microbiol. 2001;55: 453–483. doi:10.1146/annurev.micro.55.1.453

32. Gómez-Moracho T, Buendía-Abad M, Benito M, García-Palencia P, Barrios L, Bartolomé C, et al. Experimental evidence of harmful effects of Crithidia mellificae and Lotmaria passim on honey bees. Int J Parasitol. 2020;50: 1117– 1124. doi:10.1016/j.ijpara.2020.06.009

33. Buendía-Abad M, Higes M, Martín-Hernández R, Barrios L, Meana A, Fernández Fernández A, et al. Workflow of Lotmaria passim isolation: Experimental infection with a low-passage strain causes higher honeybee mortality rates than the PRA-403 reference strain. Int J Parasitol Parasites Wildl. 2021;14: 68–74. doi:10.1016/j.ijppaw.2020.12.003

34. Higes M, Rodríguez-García C, Gómez-Moracho T, Meana A, Bartolomé C, Maside X, et al. Survival of honey bees (Apis mellifera) infected with Crithidia mellificae spheroid forms (Langridge and mcGhee: atcc ® 30254TM) in the presence of Nosema ceranae. Spanish J Agric Res. 2016;14: 2171–9292. doi:10.5424/sjar/2016143-8722

35. Martín-Hernández R, Higes M, Sagastume S, Juarranz Á, Dias-Almeida J, Budge GE, et al. Microsporidia infection impacts the host cell’s cycle and reduces host cell apoptosis. PLoS One. 2017;12: 1–17. doi:10.1371/journal.pone.0170183

36. Martín-Hernández R, Aránzazu Meana, García-Palencia P, Marín P, Botías C, Garrido-Bailón E, et al. Effect of temperature on the biotic potential of honeybee microsporidia. Appl Environ Microbiol. 2009;75: 2554–2557. doi:10.1128/AEM.02908-08

37. Higes M, García-Palencia P, Martín-Hernández R, Meana A. Experimental infection of Apis mellifera honeybees with Nosema ceranae (Microsporidia). J Invertebr Pathol. 2007;94: 211–217. doi:10.1016/j.jip.2006.11.001

38. Brooker BE. Desmosomes and hemidesmosomes in the flagellate Crithidia fasciculata. Zeitschrift für Zellforsch und Mikroskopische Anat. 1970;105: 155– 166. doi:10.1007/BF00335467

39. Jensen C, Schaub GA, Molyneux DH. The effect of Blastocrithidia triatomae (Trypanosomatidae) on the midgut of the reduviid bug Triatoma infestans. Parasitology. 1990;100: 1–9. doi:https://doi.org/10.1017/S0031182000060054

40. Woodcock HM, Lond DS. Studies on Avian Haemoprotozoa: No. III.— Observations on the Development of Trypanosoma noctuae (of the Little Owl) in Culex pipiens; with Remarks on the Other Parasites occurring. Q J Microsc Sci. 1914;57. Available: https://www.cabdirect.org/cabdirect/abstract/19142901532

41. Killick Kendrick R, Molyneux DH, Ashford RW. Leishmania in phlebotomid sandflies. I. Modifications of the flagellum associated with attachment to the mid gut and oesophageal valve of the sandfly. Proc R Soc London - Biol Sci. 1974;187: 409–419. doi:10.1098/rspb.1974.0085

42. Brooks AS. Ultrastructure of the Flagellar Attachment Site in Three Species of Trypanosomatids. Trans Am Microsc Soc. 1978;97: 287. doi:10.2307/3225982

43. Vickerman K, Tetley L. Flagellar Surfaces of Parasitic Protozoa and Their Role in Attachment. Ciliary Flagellar Membr. 1990; 267–304. doi:10.1007/978-1-4613-0515-6_11

44. Molyneux DH, Killick Kendrick R, Ashford RW. Leishmania in phlebotomid sandflies. III. The ultrastructure of Leishmania mexicana amazonensis in the midgut and pharynx of Lutzomyia longipalpis. Proc R Soc London - Biol Sci. 1975;190: 341–357. doi:10.1098/rspb.1975.0098

45. Brooker BE. Flagellar adhesion of Crithidia fasciculata to Millipore filters. Protoplasma. 1971;72: 19–25. doi:10.1007/BF01281007

46. Wakid MH, Bates PA. Flagellar attachment of Leishmania promastigotes to plastic film in vitro. Exp Parasitol. 2004;106: 173–178. doi:10.1016/j.exppara.2004.03.001

47. Killick-Kendrick R, Leaney AJ, Ready PD, Molyneux DH. Leishmania in phlebotomid sandflies-IV. The transmission of Leishmania mexicana amazonensis to hamsters by the bite of experimentally infected Lutzomyia longipalpis. Proc R Soc London - Biol Sci. 1977, 196: 105–115. doi: 10.1098/rspb.1977.0032

48. Adler S. Approaches to research in leishmaniasis. Sci Rep Ist Super Sanita. 1962;2: 143–150.

49. Schaub GA, Neukirchen K, Golecki J. Attachment of Blastocrithidia triatomae (trypanosomatidae) by flagellum and cell body in the midgut of the reduviid bug Triatoma infestans. Eur J Protistol. 1992;28: 322–328. doi:10.1016/S0932-4739(11)80239-7

50. Wheeler RJ, Gluenz E, Gull K. The cell cycle of Leishmania: Morphogenetic events and their implications for parasite biology. Mol Microbiol. 2011;79: 647–662. doi:10.1111/j.1365-2958.2010.07479.x

51. Brooker BE. Flagellar attachment and detachment of Crithidia fasciculata to the gut wall of Anopheles gambiae. Protoplasma. 1971;73: 191–202. doi:10.1007/BF01275594

52. Bonilla-Rosso G, Engel P. Functional roles and metabolic niches in the honey bee gut microbiota. Curr Opin Microbiol. 2018;43: 69–76. doi:https://doi.org/10.1016/j.mib.2017.12.009

53. Raymann K, Moran NA. The role of the gut microbiome in health and disease of adult honey bee workers. Curr Opin Insect Sci. 2018;26: 97–104. doi:10.1016/j.cois.2018.02.012

54. Eduardo Serrão J, Gonçalves Santos C. Histology of the Ileum in Bees (Hymenoptera, Apoidea). Braz J morphol Sci. 2006;23: 3–4.

55. Stell I. Understanding bee anatomy: a full colour guide. Catford Press; 2012.

56. Aupperle H, Genersch E. Diagnostic colour atlas of bee pathology. Bad Kissingen: Laborklin; 2016.

57. Schaub GA. Pathogenicity of trypanosomatids on insects. Parasitol Today. 1994;10: 463–468.

58. Molyneux DH, Ashford RW. Observations on a trypanosomatid flagellate in a flea, Peromyscopsylla silvatica spectabilis. Ann Parasitol Hum Comp. 1975;50: 265–274. doi:10.1051/parasite/1975503265

59. Ramsay BYJA. Excretion by the Malpighian Tubules of the Stick Insect, Dixippus morosus (Orthoptera, Phasmidae): Amino Acids, Sugars and Urea. J Exp Biol. 1958;35: 871–891.

60. Marcilla A, Martin-Jaular L, Trelis M, de Menezes-Neto A, Osuna A, Bernal D, et al. Extracellular vesicles in parasitic diseases. J Extracell Vesicles. 2014;3. doi:10.3402/jev.v3.25040

61. Schmid-Hempel P, Schmid-Hempel R. Transmission of a pathogen in Bombus terrestris, with a note on division of labour in social insects. Behav Ecol Sociobiol. 1993;33: 319–327. doi:10.1007/BF00172930

62. Durrer S, Schmid-Hempel P. Shared use of flowers leads to horizontal pathogen transmission. Proc R Soc B Biol Sci. 1994;258: 299–302. doi:10.1098/rspb.1994.0176

63. Adler LS, Michaud KM, Ellner SP, Mcart SH, Philip C, Irwin RE. Disease where you dine: Plant species and floral traits associated with pathogen transmission in bumble bees. Ecology. 2018;99: 2535–2545. doi:10.1002/ecy.2503.

64. Arismendi N, Castro MP, Vargas M, Zapata C, Riveros G. The trypanosome Lotmaria passim prevails in honey bees of different ages and stages of development. J Apic Res. 2020;0: 1–7. doi:10.1080/00218839.2020.1828239

65. Pimenta PFP, Turco SJ, McConville MJ, Lawyer PG, Perkins P V., Sacks DL. Stage-specific adhesion of Leishmania promastigotes to the sandfly midgut. Science (80-). 1992;256: 1812–1815. doi:10.1126/science.1615326

